# Generating synthetic data with a mechanism-based Critical Illness Digital Twin: Demonstration for Post Traumatic Acute Respiratory Distress Syndrome

**DOI:** 10.1101/2022.11.22.517524

**Authors:** Chase Cockrell, Seth Schobel-McHugh, Felipe Lisboa, Yoram Vodovotz, Gary An

## Abstract

Machine learning (ML) and Artificial Intelligence (AI) approaches are increasingly applied to predicting the development of sepsis and multiple organ failure. While there has been success in demonstrating the clinical utility of such systems in terms of affecting various outcomes, there are fundamental challenges to the ML/AI approach in terms of improving the actual predictive performance and future robustness of such systems. Given that one of the primary proposed avenues for improving algorithmic performance is the addition of molecular/biomarker/genetic features to the data used to train these systems, the overall sparsity of such available data suggests the need to generate synthetic data to aid in training, as has been the case in numerous other ML/AI tasks, such as image recognition/generation and text analysis/generation. We propose the need to generate synthetic molecular/mediator time series data coincides with the advent of the concept of medical digital twins, specifically related to interpretations of medical digital twins that hew closely to the original description and use of industrial digital twins, which involve simulating multiple individual twins from a common computational model specification. Herein we present an example of generating synthetic time series data of a panel of pro- and anti-inflammatory cytokines using the Critical Illness Digital Twin (CIDT) regarding the development of post-traumatic acute respiratory distress syndrome.

## 1.0 Introduction

Predicting the course of sepsis and multiple organ failure remains a challenge. More recently machine learning (ML) methods have been applied to predict the onset of sepsis, such as the TREWS (Targeted Real-time Early Warning System) algorithm, which was able to impact clinical care by improving outcomes, primarily via the earlier administration of antibiotics [1, 2]. However, despite its promise, there are known fundamental and intrinsic limitations to ML, primarily related to challenges in generalizing to new use-contexts/cases and an inevitable performance degradation over time due to *data drift* [3-6]. In short, data drift refers to an evolution or change in the factors and processes that generate the observations/data upon which the ML algorithm operates. An example of contextual data drift (i.e., accounting for differential practices, standards, or epidemiological risks at various medical centers) would be an attempt to apply a ML algorithm trained at a particular series of institutions to a new institution with a known difference in type of microbial nosocomial infections; an example of temporal data drift would be the adoption of a new care pathway, diagnostic test, or treatment algorithm. While increasing the breadth of patient data contexts (e.g. new and varied practice sites) available for model training is the most common proposed response to addressing contextual generalizability (an approach that was effectively applied in the TREWS studies), there is no currently accepted method to account for temporal data drift other than retraining. While many have consigned themselves to the perpetual need to retrain these systems (and have developed processes that will help accomplish this while minimizing significant clinical impact) [7], there is an alternative approach drawn from industrial/technological fields that can potentially enhance the efficiency and effectiveness of utilizing existing data to increase generalizable prediction. This approach is the concept of a *digital twin*, which refers to an individualized computational representation from within a specific class of objects or type of industrial process. This term is derived from industrial/material applications and, based on the original industrial definition, a digital twin consists of three components [8]:

1. A data structure for the real-world system,
2. A computational representation of some process that links data points together to form dynamics (a *simulation*), and
3. A link to the real-world system that feeds data back into the data-propagation/generation process to produce an individualized trajectory for the specific object/process being twinned.

These three components allow the digital twin to predict/forecast (in an individualized/personalized fashion) the future history of the specific twinned objects and identify specific failure points (e.g., in a hypothetical biomedical application, *when/if* the patients will develop ARDS). Further, assessment of a digital twin that follows the classical description (e.g. it can be *simulated*) can help define potential corrective actions to be taken in real-time (e.g., *how* to prevent the patient from developing ARDS). Central to this description of a digital twin within its original industrial context is that there is a *computational specification* for the class of twinned object, from which individual instances of digital twins can be created, simulated and used.

The concept of a “medical digital twin” [9-13] embodies the idealized goals of personalized and precision medicine: identifying an individual patient’s risk of disease, a time-horizon in which clinical consequences can manifest, and potentially offering therapeutic measures to forestall a poor outcome; however at this early stage there are several different concepts for what actually constitutes a medical digital twin [9-13]. *For our purposes, we adopt a definition that follows closely the original definition of an industrial digital twin, specifically recognizing that inherent in the concept of industrial digital twins is the ability to simulate multiple possible future behavior(s) of the twin where individual simulations arise from a common shared computational specification/model*. This set of behaviors can then be refined by the data connection between the real world and the digital twin and used to narrow the forecasted set of possible futures for the twinned individual (in a fashion analogous to the way putative hurricane paths are presented as a rolling forecasting “cone” that is updated with new data). This need to *simulate* the digital twin points to the recognition within the industrial application that data-centric means of linking together time series data alone (including ML) are generally insufficient to project future trajectories [14].

We propose that the digital twin concept be applied to the critically ill/intensive care unit population, and in this work, we aim to create a Critical Illness Digital Twin (CIDT) that can be used to augment the development of ML-models that predict sepsis and multiple organ dysfunction by enhancing their robustness, performance, and generalizability. The CIDT integrates simulation models that embody mechanistic knowledge regarding systemic inflammation following trauma with ML/Artificial Intelligence (AI) methods for calibration and parameterization to more comprehensively, realistically, and functionally encompass the heterogeneity seen in clinical populations; this capability is predicated on the fact that in order to create a digital twin of *any* individual we must be able to create a digital twin of *every* individual. The integration of these complementary methods of mechanistic simulation and ML analysis will enhance the robustness and generalizability of the ML/AI predictive models by generating bio-realistic synthetic data to supplement their training (e.g., generating populations of digital twins). This process is analogous to the ubiquitous use of synthetic training data in high impact ML/AI domains such as image or speech recognition[15], but cognizant of the unique challenges inherent to time series molecular/biomarker data. Specifically, to our knowledge, there are no published instances of attempts to generate synthetic multiplex cellular/molecular-level profiling time series data. It is also recognized that the simulation model that underlies the CIDT need not be comprehensive (which is also the case of industrial digital twins); rather, it needs to be sufficient to reliably reproduce the heterogeneity of behavior/output seen across a clinical population. In prior work, we have developed ML methods that achieve the above-described challenge [16, 17].

The underlying premise to this integrative approach to the CIDT is that humans all share some core fundamental biological processes that are conserved across individuals but may manifest in different trajectories based on genetic/epigenetic/environmental differences that affect the functional responsiveness of their cellular and molecular pathways; this exactly represents the role of a common computational specification in the digital twin concept. It is this variation of mechanistic responsiveness that contributes to the differences between how individuals respond to similar insults; this is exactly the process that turns the common specification model for a class of objects into a specific digital twin. Computationally, this manifests as different parameterizations of a common model structure used to represent the variation across a population (one such example of this concept in biomedical practice is the creation of simulated patient cohorts to computationally study differential drug kinetics in the field of Quantitative Systems Pharmacology [18, 19]; another is our use of in silico trials to evaluate and discover what mechanistic interventions, representing potential drugs, could be used to treat sepsis [20-23]). To reiterate: this approach is also consistent with the industrial concept of a digital twin, where the model structure represents the formal simulation specification of the targeted object/industrial process, and a specific “twin” is created by personalizing the parameterization (and updating) of that simulation model. For purposes of using the CIDT to aid in the development of a ML/AI prediction algorithm, simulating populations of CIDTs will produce a far larger and more expansive number of training trajectories than would ever be able to be collected clinically, allowing the ML/AI a fuller range of behavioral possibilities that would encompass multiple potential application/use contexts (i.e., different populations in different institutions); this process would be vital in being able to account for subpopulations that might be under-represented in existing data sets and therefore could be under-recognized/appreciated in classical ML training. As noted above, this process of synthetically creating a much larger training data set than would be possible in terms of real world data collection is analogous to the use of synthetically generated data in high success ML applications like image analysis and natural language processing [15]. An added benefit of the mechanism-based simulation component of the CIDT is the future ability to use the CIDT for therapeutic discovery and personalized treatment optimization [21, 24].

## 2.0 Methods

In this work, the underlying common computational specification model for the CIDT is the previously developed and validated Innate Immune Response Agent-based Model (IIRABM) [17, 20, 21, 25]. In this abstraction, individual cells perform actions (i.e., apoptosis, phagocytosis, or cytokine production/secretion) using rules derived from the available literature; these rules that govern cellular actions are what we term *mechanisms* for describing our models.

### 2.1 Data Source

The Uniform Services University/Walter Reed National Medical Military Center provided data on 199 trauma patients, 92 of which developed ARDS at some point during the course of their hospitalization, matched with 107 controls that did not develop ARDS. Data elements that were used to calibrate the model include two primary types of elements: 1) vital signs/laboratory observables necessary to determine SOFA score (system-level phenotype); for the respiratory compartment, this consists of the partial pressure of oxygen, complete information regarding respiratory support, and blood oxygen saturation, representing aggregate organ function; and 2) time-series blood-serum cytokine profiles consisting of IL-1β, IL-1ra, IL-6, IL-4, IL-8, IL-10, G-CSF, IFNγ, and TNFα, sampled periodically for the duration of the patient’s hospitalization. To demonstrate our methodology for generating synthetic data with the CIDT, we use simulations of the IIRABM to identify parameterizations (which represent varied individuals responding to varied insults) that cannot be falsified by the available data. Using the nested GA parameter discovery and AL parameterization boundary identification noted below we determined three sets of model parameterizations: 1) patients that will certainly develop ARDS; 2) patients that certainly will not develop ARDS; and 3) patients that have a probability greater than zero, but less than one, of developing ARDS.

### 2.2 Calibration Procedure

The CIDT is calibrated and validated in a two-step procedure. First, Genetic Algorithms (GA) are used to discover the model parameterizations that best fit the full range of data. Next, an Artificial Neural Network (ANN) is trained using Active Learning (AL) to take model parameterizations as input and classify if they are valid (whether or not they are invalidated by available data); the AL procedure then allows for the discovery of boundaries and parameter rangers around GA-discovered parameterizations. A more detailed description of these processes are noted below.

#### 2.2.1 Genetic Algorithms for Parameter Discovery

The GA [26-28] is a population-based metaheuristic optimization algorithm that is inspired by biological evolution. We have previously utilized GA both to develop personalized therapeutic strategies [21] and to calibrate a model of the human innate immune response, the IIRABM, to a burn data set, simultaneously discovering novel rules regions associated model parameterizations which led to biologically plausible output [17]. Ultimately, the goal of the GA is to evolve the model’s rule set such that the solution replicates not only what is seen experimentally, but also its associated variance. The form of the fitness function we will use is:

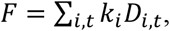

where

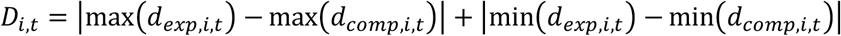

and where max(*d*_*exp,i,t*_) represents the maximum value (from all subjects at that time point) for the experimental measurement of data element *i* at time *t*; max (*d*_*comp,i,t*_,) represents the maximum value of computational model output (from all stochastic replicate simulations) for data element *i* at time *t*; and the constants *k*_*i*_ allow for varying weights to be placed on individual data elements (i.e., we would not expect the raw difference in a protein concentration to be on the same scale as differential cell populations over time). By summing the differences between the maximum and minimum values of the computational model output and experimentally collected data, this instantiation of the GA seeks to find model parameterizations that generate a range of data which most closely matches the range of data experimentally and does so by operating on the model parameterization such that, when instantiated in the model, lead it to minimize the above fitness function. We note that initially we will assign equal weights to each cytokine concentration, vital sign, or laboratory observable to be matched. As the process continues, weights may be adjusted to bias the algorithm to better fit certain aspects of the data, e.g., if the GA is slow to converge to an adequate solution that explains IL10 dynamics, this data element would be assigned a greater weight to increase its contribution to model training.

We note we are employing GA in a non-standard fashion, where rather than seeking a specific optimal parameterization of the model we are using the process of convergence of the GA to identify an ensemble of valid parameterizations. To construct this ensemble, we add an ensemble retainment criterion to the GA procedure, the rational for which is as follows: we recognize that any putative parameterization which generates cytokine trajectories that always lie within the cilnically observed range cannot be invalidated by the data, and are therefore biologically plausible; thus, these parameterizations should be retained for inclusion into the final ensemble. As the goal of the fitness function is to obtain maximum coverage over the clinical data range, some of these viable parameterizations may otherwise be lost as the population evolves.

#### 2.2.2 Active Learning to Determine Boundaries of GA-Derived Candidate Parameterizations

Active learning (AL) is a sub-field of machine learning (ML) which focusses on finding the optimal selection of training data to be used to train a ML or statistical model [29]. AL can be used for classification [30, 31] or regression [32, 33]. AL is an ideal (in terms of total cost) technique for modeling problems in which there is a large amount of unlabeled data and manually labelling that data is expensive. In these circumstances (specifically the costly data labelling) *AL provides the most generalizable and accurate model for the cheapest cost*. The problem of combinatorial complexity in the selection of model parameters is well-established in the computational/biological modeling communities [34-38]. In previous work [25], we utilized high-performance computing to demonstrate the need for comprehensive “data coverage” among possible model states as well as the importance of internal parameter variation (as compared to model structure) to capture the full range of biological heterogeneity seen clinically. AL [16] rendered this problem computationally tractable by reducing the number of simulations require to characterize the search space by ∼95% when compared to a brute-force exploration. Additionally, we demonstrated that more complex models with a larger number of variables may expect further improvements in efficiency [16].

The output of the AL workflow is an artificial neural network (ANN) classifier, for which the classes are biologically plausible (cannot be invalidated by the data) or not. After the application of the GA procedure, we will have an ensemble of biologically plausible model parameterizations. Each of these parameterizations represents a single point in a high-dimensional space. *AL will allow us to determine the biologically plausible parameter ranges around those single points in a computationally efficient manner*. We will begin the AL procedure by defining initial search boundaries, and we will discretize the search space lying within those boundaries, creating a set of putative model parameterizations with unknown classification. Each parameterization contained within the discretized search space will then be fed into the trained ANN, and have its associated class predicted. We then posit the existence of some function,

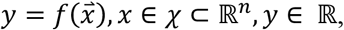

which accepts as input a model parameterization and predicts the associated class, and that this function can be approximated given input data from the training set:

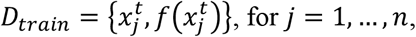

where 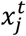 represents a single parameterization used to train the ANN. The ANN model uses a binary cross-entropy [39] loss function, in which the loss is given by:

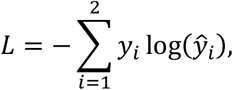

where *y*_*i*_ is the ground truth value and *ŷ*_*i*_ is the ANN-approximated score. To generate the initial training set, 100 putative parameterizations will be selected from the discretized set. The IIRABM simulation then uses these putative parameterizations runs a fixed number of stochastic replicates of the input points to determine class membership. This information is then used to train the ANN classifier. The algorithm then ranks the remaining unlabeled parameterizations by class-membership uncertainty:

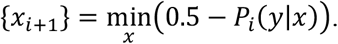

Those parameterizations whose class (biologically plausible or not) are most uncertain in the current ANN classifier model are then selected for labeling and the process repeats until a stopping criterion is reached.

## 3.0 Results

In Figures 1 and 2, we demonstrate the preliminary generation of synthetic data for the lung compartment of the CIDT. In Figure 1, Panel A, colored points are the clinically collected data, with red points representing patients that developed ARDS sometime between days 6 and 10 and blue points representing patients that did not. The dotted lines (shown only in Panel A, TNFα for clarity) indicate representative clinical trajectories, which are sparse and typically only contain 3-4 measurements. The non-ARDS clinical trajectory, for example, only contains measurements at t=0, 3, and 7 days post-hospitalization, leaving significant uncertainty as to the patient state in intermediate times. This feature is present for nearly all mediators and patients in the currently available data. The shading indicates the boundaries of the model behavior space for the parameterizations that generate ARDS (red) or not (blue), with significant overlap between the two spaces. In Panel B, we show actual simulated trajectories for blood-serum concentration of TNFα, noting the similar, but much richer, dynamics compared to the clinical trajectories. The two patients whose trajectories are represented by the red lines ultimately develop ARDS, while the patient represented by the blue line does not. In Figure 2, we show the clinical data, simulation boundaries, and simulated trajectories for blood-serum GCSF and IL-10 in Panels A and B. In Panel C, we show the oxygen deficit trajectories, an inverse-measure of patient health, for 500 synthetic patients each from the parameterizations of the model that develop ARDS (red) and those that to not (blue), with SOFA scores correlated to the clinical data, using the Berlin definition of ARDS, [40] in which mild ARDS corresponds to a Respiratory SOFA score of 2, moderate ARDS corresponds to a Respiratory SOFA score of 3, and severe ARDS corresponds to a Respiratory SOFA score of 4. It should also be noted that different clinical patients may only manifest some of the mediator variations that can distinguish between ARDS and non-ARDS; therefore, this heterogeneity of response can only be identified across a population of parameterized simulations that can generate the entire set of mediators in concert.

**Figure 1:**
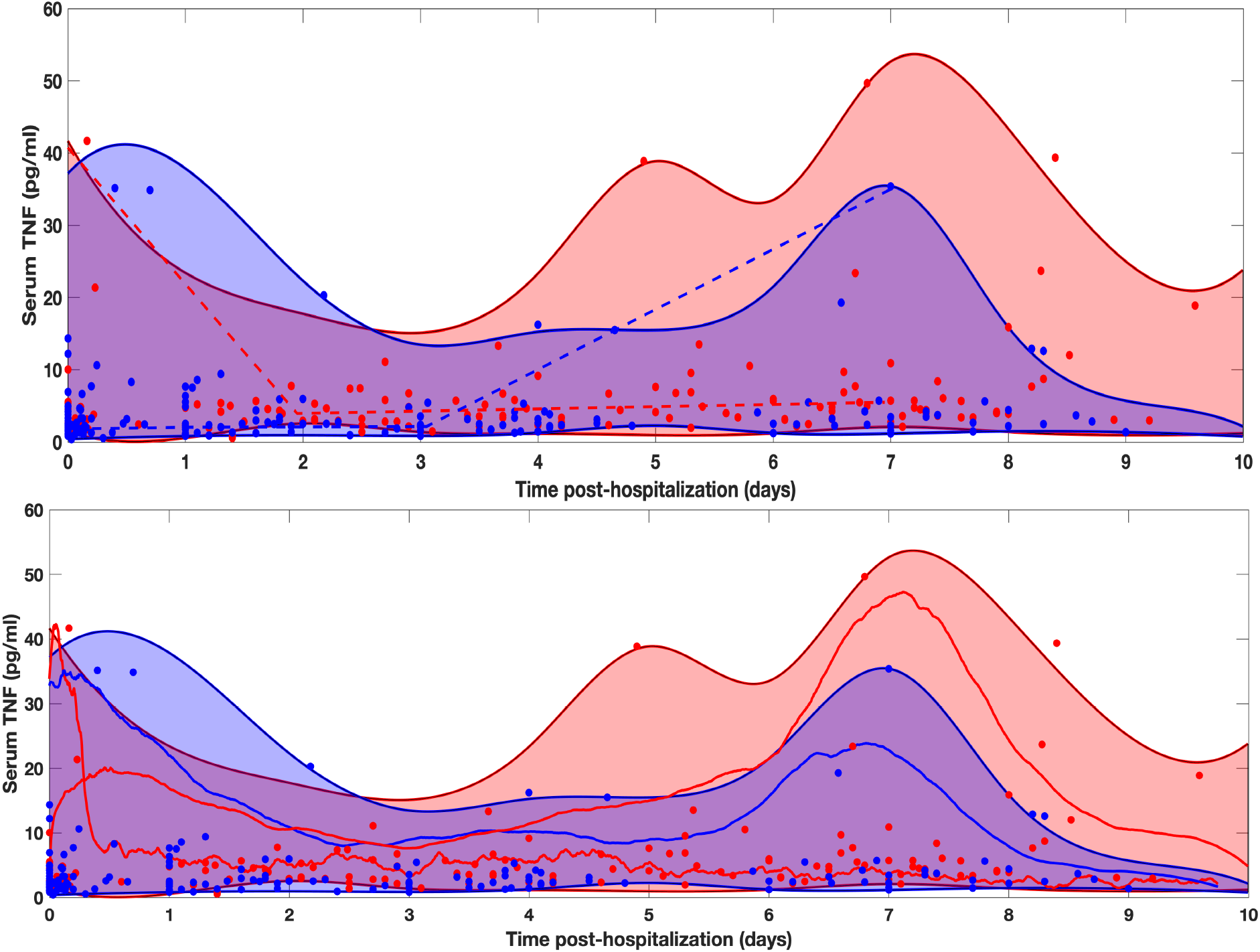
Comparison of Experimental and Simulated Protein Concentration Trajectories. In Panel A, we show both simulated and clinically collected blood-serum cytokine measurements for TNFα. Distinct points are the clinically collected data, with red points representing patients that developed ARDS sometime between days 7 and blue points representing patients that did not. The dotted lines indicate representative clinical trajectories. The shading indicates the boundaries of the model trajectory space for the parameterizations that generate ARDS (red) or not (blue). Note significant overlap between the two spaces given the overlap of data points. However, the key point is that differential parameterizations are able to identify clear regions that are unique to each group. In Panel B, the three solid lines (two red lines eventually developing ARDS and the blue line not) show actual simulated trajectories of TNFα blood-serum concentrations generated by the lung compartment of the CIDT.

**Figure 2:**
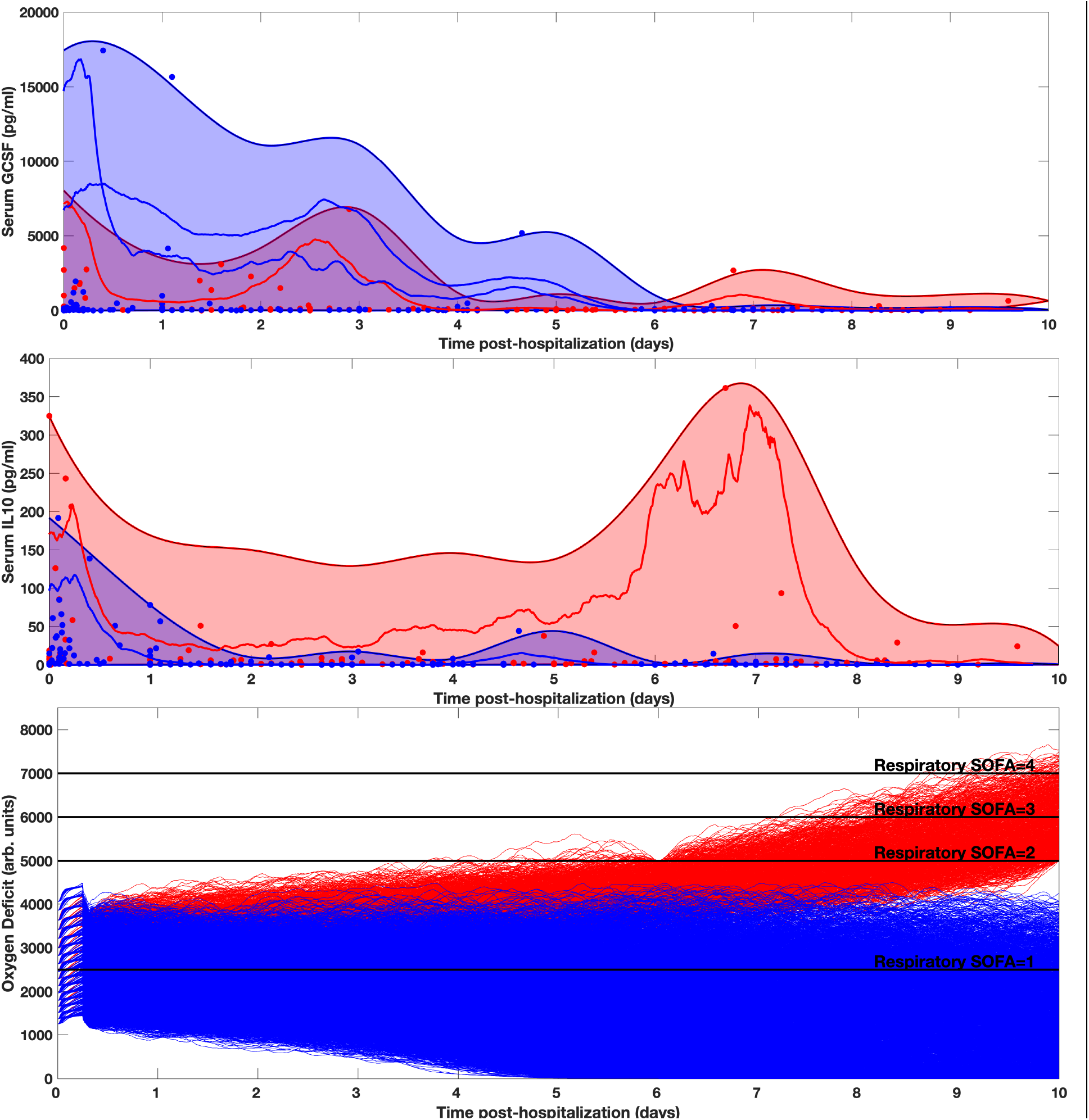
Correspondence Between Cytokine Profile and Clinical State. As in Figure 3, distinct points are the clinically collected data, with red points representing patients that developed ARDS sometime between days 7 and blue points representing patients that did not. The shading indicates the boundaries of the model behavior space for the parameterizations that generate ARDS (red) or not (blue). Simulated trajectories are shown for GCSF (Panel A) and IL10 (Panel B). In Panel C, we show the oxygen-deficit trajectories of simulated patients and their corresponding respiratory SOFA scores as they develop ARDS (increasing oxygen deficit) or progress towards successful healing of the lung injury.

## 4.0 Discussion

While sepsis prediction algorithms, such as TREWS, have been demonstrated to improve outcome, presumed in the case of TREWS to be due primarily to the earlier administration of antibiotics, there is ongoing interest in being able to augment the physiological features used by these early warning systems by adding more detailed, biologically informative features that would help discriminate between the disease trajectories of individuals. This desire is consistent with the general trend towards more precise and personalized therapy. With respect to forecasting the potential progression to ARDS, numerous ML predictive algorithms that operate on data from the electronic health record [41, 42] (also see Ref [43] for a review) have been augmented with radiographic [44-46] and respiratory/ventilator waveform [47-49] data. As in other fields, such as precision oncology, there is also interest in being able to extend the feature set to include biomarkers and genetic/epigenetic information (see Ref [50] for a review). It is in this latter group that the ability to generate synthetic time series data becomes important using a mechanism-based Digital Twin. The reason for this is the Curse of Dimensionality [51] as related to the logistical challenges of collecting sufficient samples in a clinical setting: in short, the addition of numerous features/variables to a predictive data set leads to a combinatorial explosion of the number of samples needed to identify/infer a relationship between those features and the desired goal. Since the explicit goal of a clinical forecasting/prediction algorithm is to define the risk probability for an *individual* patient/trajectory, representing and generating these potential trajectories requires deconstruction of the aggregating assumptions used for creating statistical models and in standard calibration/parametrization methods for dynamic computational models. The focus on simulating individual trajectories is inherent to the concept of digital twins, which is why we believe that the implementation of this concept in a biomedical context using mechanism-based simulation models as the underlying digital twin specification presents a possible solution to the Curse of Dimensionality, and, by extension, the challenge of dealing with temporal data drift for the implementation and maintenance of AL/ML predictive algorithms (since the wider space of possibility, and therefore generalizability, is captured by the creation of synthetic data). Inherent to this application of mechanism-based computer simulations is the calibration phase, where instead of fitting to a mean of highly variable and often overlapping data points, the simulated trajectories produce a space that encompasses the full range of the available data (see Figures 1 and 2); we present a particular workflow that accomplishes this goal [16, 17] and hope that others will develop analogous approaches. While there are clear challenges to this approach in the biomedical arena, with many overlaps with the underlying issues in translating the industrial digital twin concept to biomedicine (specifically with respect to the development of “trustworthy” mechanism-based simulation specifications [52]), this use of medical digital twins for the purpose of generating synthetic time series for the training of ML/AI forecasting algorithms represents a path forward to what are otherwise intractable limitations regarding the utility of pure ML/AI systems to forecast sepsis and multiple organ failure.

## 5.0 Summary

Synthetic data is proving to be invaluable in the training of modern ML/AI systems. There are specific challenges to generating synthetic time series data of molecular features of biological systems; these challenges include but are not limited to data sparsity, the Curse of Dimensionality and over-fitting. We propose that these challenges can be overcome using sufficiently complex mechanism-based models calibrated in a fashion that encompasses all available training data by generating distinct system trajectories. The further development of these models and methods can provide a means of generating synthetic time series data that can increase the robustness and generalizability of ML/AI systems applied to clinical scenarios where molecular/biomarker/gene expression time series data is of interest for the prediction/forecasting of disease progression and the discovery/testing of novel drug targets.

